# Mapping quantitative trait loci underlying function-valued traits using functional principal component analysis and multi-trait mapping

**DOI:** 10.1101/025577

**Authors:** Il-Youp Kwak, Candace R. Moore, Edgar P. Spalding, Karl W. Broman

**Affiliations:** Departments of Statistics, University of Wisconsin–Madison, Madison, Wisconsin 53706; Departments of Botany, University of Wisconsin–Madison, Madison, Wisconsin 53706; Departments of Biostatistics and Medical Informatics, University of Wisconsin–Madison, Madison, Wisconsin 53706

**Keywords:** QTL, function-valued traits, model selection, growth curves, multivariate analysis

## Abstract

We previously proposed a simple regression-based method to map quantitative trait loci underlying function-valued phenotypes. In order to better handle the case of noisy phenotype measurements and accommodate the correlation structure among time points, we propose an alternative approach that maintains much of the simplicity and speed of the regression-based method. We overcome noisy measurements by replacing the observed data with a smooth approximation. We then apply functional principal component analysis, replacing the smoothed phenotype data with a small number of principal components. Quantitative trait locus mapping is applied to these dimension-reduced data, either with a multi-trait method or by considering the traits individually and then taking the average or maximum LOD score across traits. We apply these approaches to root gravitropism data on Arabidopsis recombinant inbred lines and further investigate their performance in computer simulations. Our methods have been implemented in the R package, funqtl.

## Introduction

Technology developments have enabled the automated acquisition of numerous phenotypes, included function-valued traits, such as phenotypes measured over time. High-dimensional phenotype data are increasingly considered as part of efforts to map the genetic loci (quantitative trait loci, QTL) that influence quantitative traits.

Numerous methods are available for QTL mapping with function-valued traits. Ma *et al.* (2002) considered parametric models such as the logistic growth model, 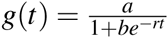. The high-dimensional phenotype is reduced to a few parameters. This works well if the parametric model is approximately correct but in many cases the correction functional form is not clear. Yang *et al.* (2009) proposed a non-parametric functional QTL mapping method, with a selected number of basis functions to fit the function-valued phenotype. Min *et al.* (2011), Sillanpää *et al.* (2012), and Li and Sillanpää (2013) extended this method for multiple-QTL models. Min *et al.* (2011) used Markov chain Monte Carlo (MCMC), Sillanpää *et al.* (2012) used hierarchical modeling, and Li and Sillanpää (2013) used a multivariate regression method. Xiong *et al.* (2011) proposed an additional non-parametric functional mapping method based on estimating equations. An important barrier to these methods is the long computation time required for the analysis.

In Kwak *et al.* (2014), we proposed two simple regression-based methods to map quantitative trait loci underlying function-valued phenotypes, based on the results from the individual analysis of the phenotypes at each time point. With the SLOD score, we take the average of the LOD scores across time points, and with the MLOD score, we take the maximum. These approaches are fast to compute, work well when the trait data are smooth, and provide results that are easily interpreted. Another important advantage is the ability to consider multiple QTL, which can improve power and enable the separation of linked QTL. However, the approaches do not work as well when the trait data are not smooth, and they do not take account of the correlation among time points.

In the present paper, we describe methods to overcome these weaknesses. First, we replace the observed trait data with a smooth approximation. Second, we apply functional principal component analysis (PCA) as a dimension-reduction technique, and replace the smoothed phenotype data with a small number of principal components.

QTL analysis is then performed on these dimension-reduced data. We consider either the multivariate QTL mapping method of Knott and Haley (2000), or use the SLOD or MLOD scores, as in Kwak *et al.* (2014). These methods all have analogs for multiple-QTL models, by extending the penalized LOD scores of Broman and Speed (2002) and Manichaikul *et al.* (2009).

We illustrate these methods by application to the root gravitropism data of Moore *et al.* (2013), measured by automated image analysis over a time course of 8 hr across a population of *Arabidopsis thaliana* recombinant inbred lines (RIL). We further investigate the performance of these approaches in computer simulations.

## Methods

We will focus on the case of recombinant inbred lines, with genotypes AA or BB. Consider *n* lines, with function-valued phenotypes measured at *T* discrete time points (*t*_1_, ⋯, *t_T_*).

### Smoothing

We first smooth the phenotype data for each individual. Let *y_i_(t_j_)* denote the observed phenotype for individual *i* at time *t_j_*. We assume underlying smooth curves *x_i_(t*), with *y_i_(t_j_) = x_i_(t_j_*) + *e_i_(t_j_*).

We approximate the functional form of *x_i_(t)* as 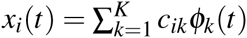 for a set of basis functions *ϕ*_1_ *(t*), ⋯, *ϕ_K_(t*), where the number of basis functions, *K*, is generally much smaller than the number of time points, *T*. There are many possible choices of basis functions. We are using B-splines (Ramsay and Silverman 2005, pp. 49-53). Xiong *et al.* (2011) and Li and Sillanpää (2013) also used B-splnes for QTL analysis with functional traits.

Define the *K* by *T* matrix Φ, with Φ*_kj_ = ϕ_k_(t_j_*), for *k* = 1, …,*K* and *j* = 1,…, *T*. We estimate the coefficients, *c_ik_* by minimizing the least squares criterion (Ramsay and Silverman 2005):

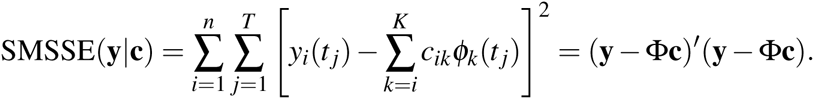

This gives the solution 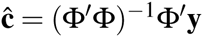. We then obtain the smoothed phenotypes 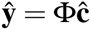, which are used in all subsequent analyses.

A key issue is the choice of the number of basis functions. We use 10-fold cross-validation and choose the number of basis functions that minimizes the estimated sum of squared errors.

### Functional principal component analysis

Having replaced the phenotype data with a smooth approximation, 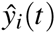, we use functional principal component analysis (Ramsay and Silverman 2005) to reduce the dimensionality with little loss of information.

In functional PCA, we seek a sequence of orthonormal functions, *ψ_j_(t*), that take the role of principal components but for functional data. By orthonormal, we mean 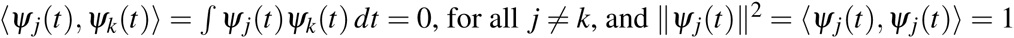, for all *j* ≠ k, and 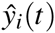 for all *j.*

With our smoothed functional data, 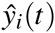, we first find the function *ψ_1_(t*) that maximizes 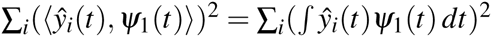, subject to the constraint 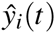. The computations make use of the B-spline basis representation of the 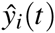.

Conceptually, the procedure then proceeds inductively. Having identified *ψ*_1_,…, *ψ_j_*, we choose *ψ_j_*_+1_ that maximizes 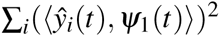 subject to the constraint that *ψ*_1_,…, *ψ_j_*_+1_ are orthonormal.

We focus on a small number, *p*, of functional PCs that explain 99% of the data variation and consider the coefficients 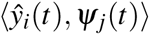, as derived traits.

### Single-QTL analysis

Having smoothed and dimension-reduced the phenotype data, we then use the *p* principal components as derived traits for QTL analysis, using one of three methods. First, we apply the multivariate QTL mapping method of Knott and Haley (2000). Our second and third approaches are to analyze the *p* derived traits individually and take the average or maximum LOD score, respectively, at each putative QTL position, as in Kwak *et al.* (2014).

**HKLOD score:** In the method of Knott and Haley (2000), we take *n* × *p* matrix of derived phenotypes and scan the genome, and at each position, *λ*, we fit a multivariate regression model with a single QTL. The basic model is *Y = XB* + *E*, where *Y* is the *n* × *p* matrix of phenotypes, *X* is an *n* × 2 matrix of QTL genotype probabilities, and *B* is a 2 × *p* matrix of QTL effects. The rows of the *n* × *p* matrix of errors, *E*, are assumed to be independent and identically distributed draws from a multivariate normal distribution.

The maximum likelihood estimate of the coefficients is 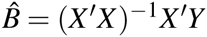, the same as if the traits were analyzed separately. A key component of the likelihood is the matrix of sums of squares and cross-products of residuals, RSS = *(Y* − *XB)′(Y* − *XB),* and particularly its determinant, |RSS|. The log_10_ likelihood ratio comparing the model with a single QTL at position *λ*, to the null model of no QTL, is

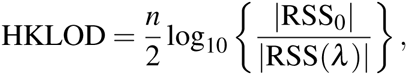

where |RSS_0_| is for the null model with no QTL and |RSS(*λ*) is for the model with a single QTL at *λ*.

**SL and ML scores:** As further approaches, we apply the method of Kwak *et al.* (2014) to the *p* derived traits. We perform a genome scan by Haley-Knott regression (Haley and Knott 1992) with each trait separately, giving the *LOD_j_ (λ)* for trait *j* at position *λ*.

Our second criterion, is to take the average LOD score, across traits: 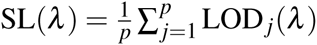. We call this the SL score, to distinguish it from SLOD of Kwak *et al.* (2014), calculated using the original phenotype data.

Our third criterion is to take the maximum LOD score, ML(*λ*) = max*_j_* LOD*_j_* (*λ*). We call this the ML score, to distinguish it from the MLOD score of Kwak *et al.* (2014).

### Multiple-QTL analysis

As in Kwak *et al.* (2014), we use the penalized LOD score criterion on Broman and Speed (2002) to extend each of the LOD-type statistics defined above, for use with multiple-QTL models. The penalized LOD score is pLOD(*γ*) = LOD(*γ*) − T|*ɣ*|, where *ɣ* denotes a multiple-QTL model with strictly additive QTL, and |*ɣ*| is the number of QTL in the model *γ*. *T* is a penalty on model size, chosen as the 1 − *α* quantile of the genome-wide maximum LOD score under the null hypothesis of no QTL, derived from a permutation test (Churchill and Doerge 1994). We may replace the LOD score in the above equation with any of the HKLOD, SL and ML scores.

To search the space of models, we use the stepwise model search algorithm of Broman and Speed (2002): we use forward selection up to a model of fixed size (e.g., 10 QTL), followed by backward elimination to the null model. The selected model 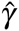 is that which maximizes the penalized LOD score criterion, among all models visited.

The selected model is of the form *Z = Qβ* + *ε*, where *Z* contains the derived traits (the coefficients from the functional principal component analysis) and *Q* contains an intercept column and genotype probabilities at the inferred QTL. The derived phenotypes, *Z*, are linearly related to the smoothed phenotypes, 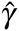, by the equation 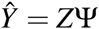, where Ψ is a matrix with (*i*, *j*)th element *ψ_i_(t_j_*). Thus we have 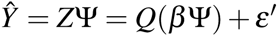, and so 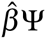 are the estimated QTL effects, translated back to the time domain (see Figure 3, below).

## Application

As an illustration of our approaches, we considered data from Moore *et al.* (2013) on gravitropism in Arabidopsis recombinant inbred lines (RIL), Cape Verde Islands (Cvi) × Landsberg erecta (Ler). For each of 162 RIL, 8–20 replicate seeds per line were germinated and then rotated 90 degrees, to change the orientation of gravity. The growth of the seedlings was captured on video, over the course of eight hours, and a number of phenotypes were derived by automated image analysis.

We focus on the angle of the root tip, in degrees, over time (averaged across replicates within an RIL), and consider only the first of two replicate data sets examined in Moore *et al.* (2013). There is genotype data at 234 markers on five chromosomes; the function-valued root tip angle trait was measured at 241 time points (every two minutes for eight hours).

The data are available at the QTL Archive, which is now part of the Mouse Phenome Database, as the *Moore1b* data set:

http://phenome.jax.org/db/q?rtn=projects/projdet&reqprojid=282

### Single-QTL analysis

We first performed genome scans with a single-QTL model by the multiple methods: SLOD and MLOD from Kwak *et al.* (2014), EE(Wald) and EE(Residual) from Xiong *et al.* (2011), and the HKLOD, SL, and ML methods described above (and using four principal components). We used a permutation test (Churchill and Doerge 1994) with 1000 permutation replicates to estimate 5% significance thresholds, which are shown in Supporting Information, Table S1.

The results are shown in Figure 1. The SLOD method (Figure 1A) gave similar results to the EE(Residual) method (Figure 1D), with significant evidence for QTL on chromosomes 1, 4, and 5. The MLOD method (Figure 1B) also showed evidence for a QTL on chromosome 3. The EE(Wald), HKLOD, and SL methods (Figure 1C, 1E, and 1F) all gave similar results, with significant evidence for QTL on each chromosome. The ML method (Figure 1G) is different, with significant evidence for QTL on chromosomes 1, 2, and 4.

**Figure 1:**
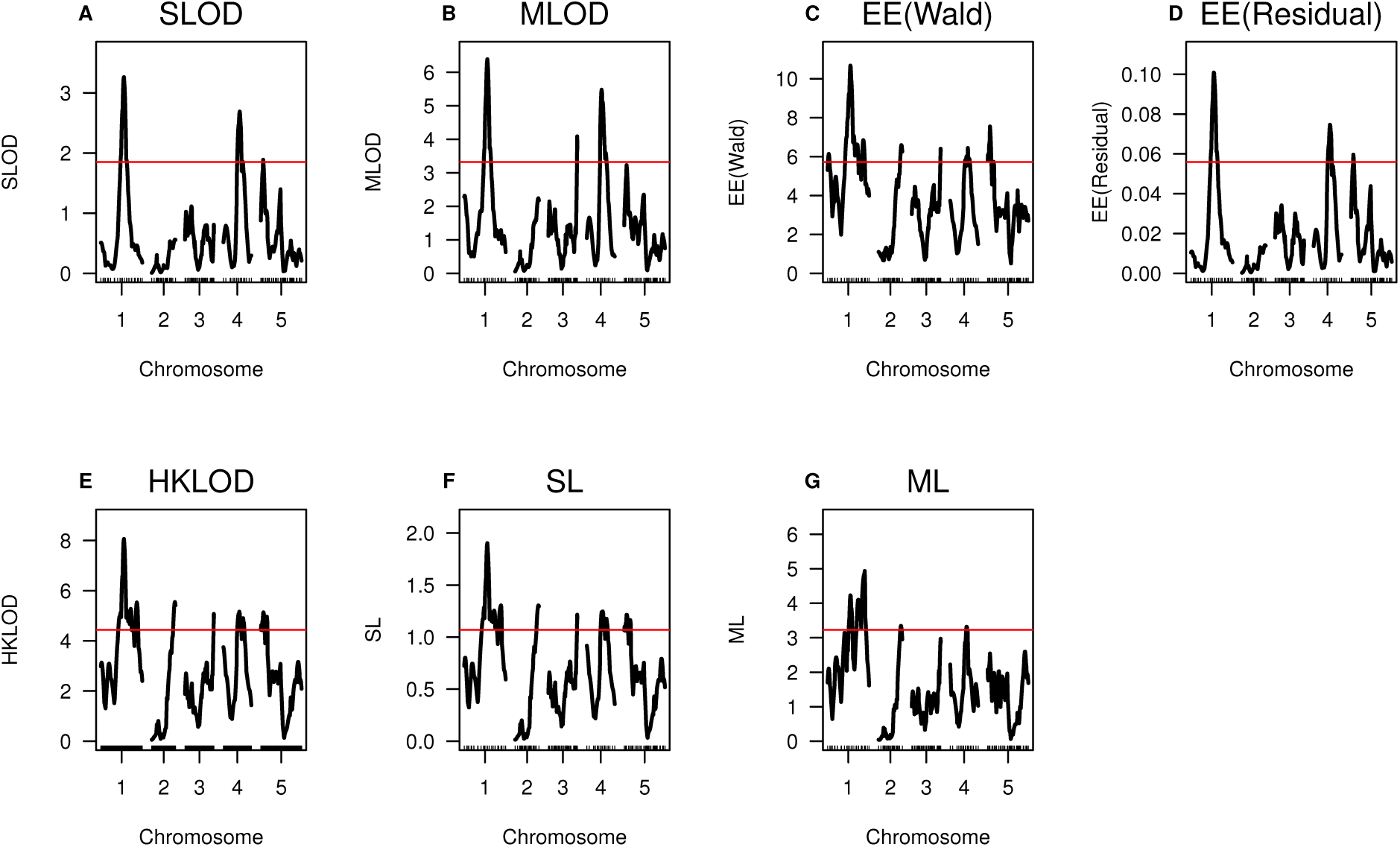
The SLOD, MLOD, EE(Wald) and EE(Residual), HKLOD, SL and ML curves for the root tip angle data. A red horizontal line indicates the calculated 5% permutation-based threshold.

### Multiple-QTL analysis

We further applied multiple-QTL analysis, extending the HKLOD, SL, and ML methods to use the penalized LOD score criterion of Broman and Speed (2002) for function-valued traits. We focused on additive QTL models, and we used the 5% permutation-based significance thresholds (Table S1) as the penalties.

The penalized-HKLOD and penalized-SL criteria each indicated a five-QTL model, with a QTL on each chromosome. The inferred positions of the QTL showed only slight differences. The penalized-SL criterion indicated a two-QTL model, with QTL on chromosomes 1 and 4.

LOD profiles for these models are displayed in Figure 2. These curves, which visualize both the evidence and localization of each QTL in the context of a multiple-QTL model, are calculated following an approach developed by Zeng *et al.* (2000): The position of each QTL was varied one at a time, and at each location for a given QTL, we derived a LOD-type score comparing the multiple-QTL model with the QTL under consideration at a particular position and the locations of all other QTL fixed, to the model with the given QTL omitted. For the SL (or ML) method, the profile is calculated for the four derived traits, individually, and then the SL (or ML) profiles are obtained by averaging (or maximizing) across traits. For the HKLOD method, the profiles are calculated using the multivariate LOD test statistic.

**Figure 2:**
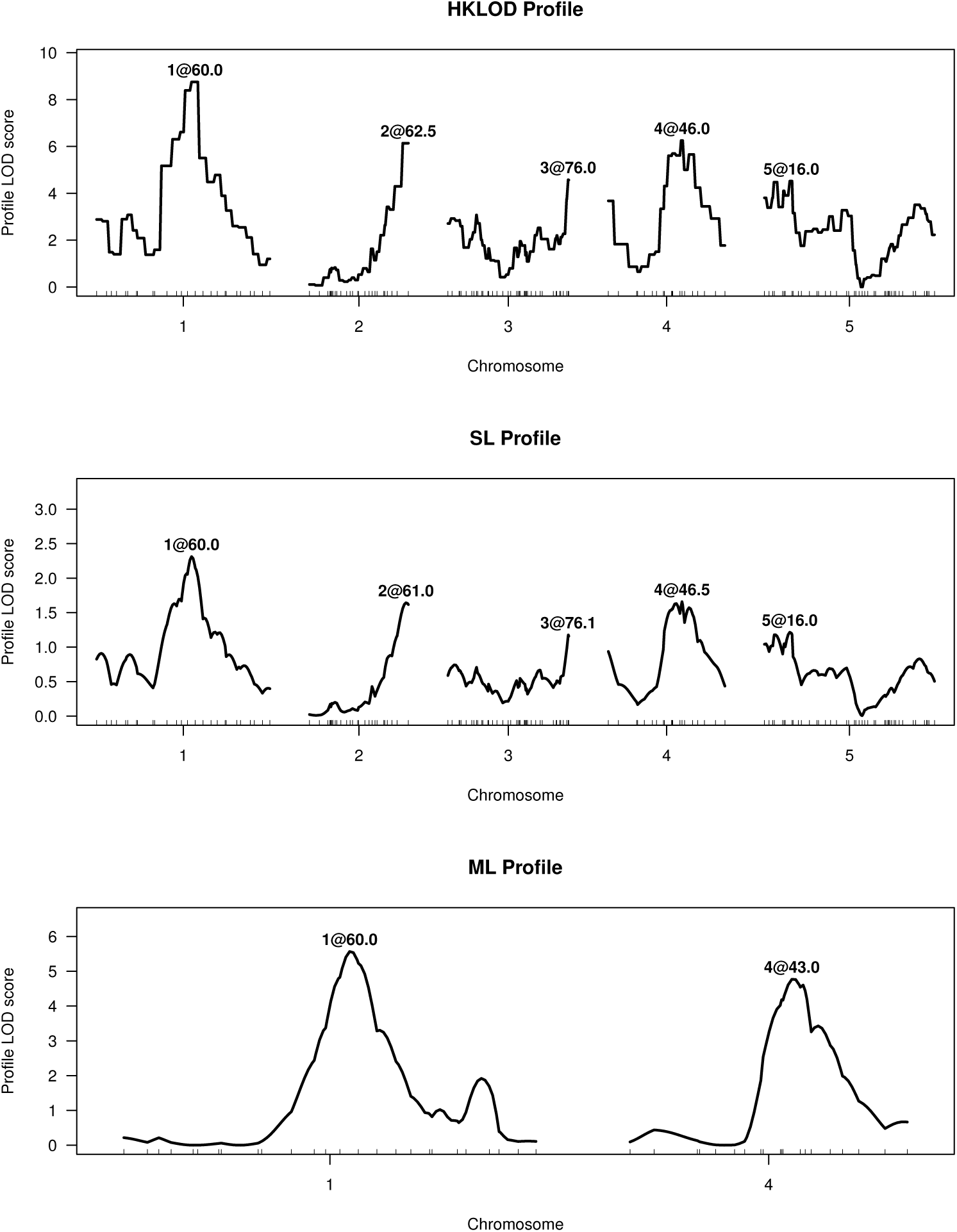
HKLOD, SL, and ML profiles for a multiple-QTL model with the root tip angle data set.

In Kwak *et al.* (2014), we applied, to these same data, the analogous multiple-QTL modeling approach, with the penalized-SLOD and penalized-MLOD criteria (that is, using the original phenotype data, without the smoothing and dimension-reduction steps). The result (see Figure 3 in Kwak *et al.* 2014) with the penalized-SLOD criterion was the same two-QTL model identified by ML (Figure 2C), while the penalized-MLOD criterion gave a three-QTL model, with a QTL on chromosome 3, similar to that inferred by single-QTL analysis with MLOD (Figure 1B).

**Figure 3:**
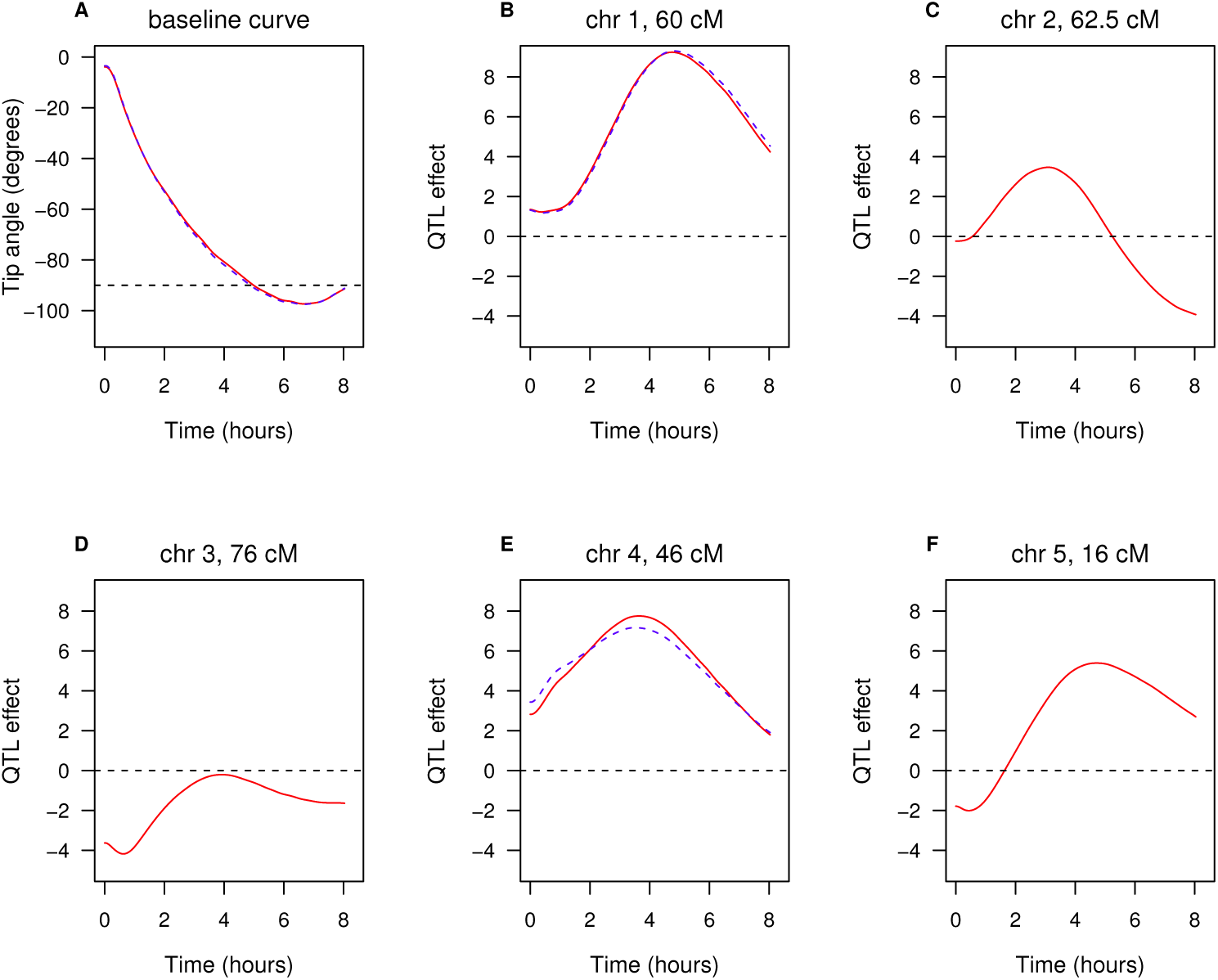
The estimated QTL effects for the root tip angle data set. The red curves are for the five-QTL model (from the penalized-HKLOD and penalized-SL criterion) and the blue dashed curves are for the two-QTL model (from the penalized-ML criterion). Positive values for the QTL effects indicate that the Cvi allele increases the tip angle phenotype.

The estimated QTL effects, translated back to the time domain, are displayed in Figure 3. The red curves are for the five-QTL model identified with the penalized-HKLOD and penalized-SL criteria. The blue dashed curves are for the two-QTL identified with the penalized-ML criterion. The estimated QTL effects in panels B–F are for the difference between the Cvi allele and the Lerallele.

The effects of the QTL on chromosomes 1 and 4 are approximately the same, whether or not the chromosome 2, 3 and 5 QTL are included in the model. The chromosome 1 QTL has greatest effect at later time points, while the chromosome 4 QTL has greatest effect earlier and over a wider interval of time. For both QTL, the Cvi allele increases the root tip angle phenotype.

In summary, the HKLOD and SL methods gave similar results. For these data, the HKLOD and SL methods indicate evidence for a QTL on each chromosome. The results suggest that these approaches, with the initial dimension-reduction via functional PCA, may have higher power to detect QTL, as the multiple-QTL analysis with SLOD and MLOD, without dimension-reduction, indicated fewer QTL.

## Simulations

In order to investigate the performance of our proposed approaches and compare them to existing methods, we performed computer simulation studies with both single-QTL models and models with multiple QTL. For the simulations with a single-QTL model, we compared the HKLOD, SL, and ML methods to the SLOD and MLOD methods of Kwak *et al.* (2014) and the estimating equation approaches of Xiong *et al.* (2011). For the simulations with multiple QTL, we omitted the methods of Xiong *et al.* (2011), as they have not yet been implemented for multiple QTL.

### Single-QTL models

To compare approaches in the context of a single QTL, we considered the simulation setting described in Yap *et al.* (2009), though exploring a range of QTL effects.

We simulated an intercross with sample sizes of 100, 200, or 400, and a single chromosome of length 100 cM, with 6 equally spaced markers and with a QTL at 32 cM. The associated phenotypes was sampled from a multivariate normal distribution with the mean curve following a logistic function, 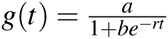. The AA genotype had *a* = 29, *b* = 7, *r* = 0.7; the AB genotype had *a* = 28.5, *b* = 6.5, *r* = 0.73; and the BB genotype had *a* = 27.5, *b* = 5, *r* = 0.75. The shape of the growth curve with this parameter was shown in Figure S3 of Kwak *et al.* (2014). The phenotype data were simulated at 10 time points.

The residual error was assumed to following multivariate normal distributions with a covariance structure *c*Σ. The constant *c* controls the overall error variance, and Σ was chosen to have one of three forms: (1) auto-regressive with *σ*^2^ = 3, *ρ* = 0.6, (2) equicorrelated with *σ*^2^ = 3, *ρ* = 0.5, or (3) an “unstructured” covariance matrix, as given in Yap *et al.* (2009) (also shown in Table S2 of Kwak *et al.* 2014)).

The parameter *c* was given a range of values, which define the percent phenotypic variance explained by the QTL (the heritability). The effect of the QTL varies with time; we took the mean heritability across time as an overall summary. For the auto-regressive and equicorrelated covariance structures, we used *c* = 1,2,3,6; for the unstructured covariance matrix, we took *c* = 0.5,1,2,3.

For each of 10,000 simulation replicates, we applied the previous SLOD and MLOD methods of Kwak *et al.* (2014), the EE(Wald) and EE(Residual) methods of Xiong *et al.* (2011), and the HKLOD, SL, and ML methods proposed here. For all seven approaches, we fit a three-parameter QTL model (that is, allowing for dominance).

The estimated power to detect the QTL as a function of heritability due to the QTL, for *n* = 100,200,400 and for the three different covariance structures, is shown in Figure 4. With the autocorrelated variance structure, all methods worked well, though HKLOD had noticeably lower power. With the equicorrelated variance structure, EE(Wald), SL and ML methods had higher power than the other four methods. The HKLOD method also worked reasonably well, but the EE(Residual), SLOD and MLOD methods had low power. In the unstructured variance setting, EE(Wald), MLOD, SL and ML methods worked better than the other three methods. EE(Residual) did not work well in this setting.

**Figure 4:**
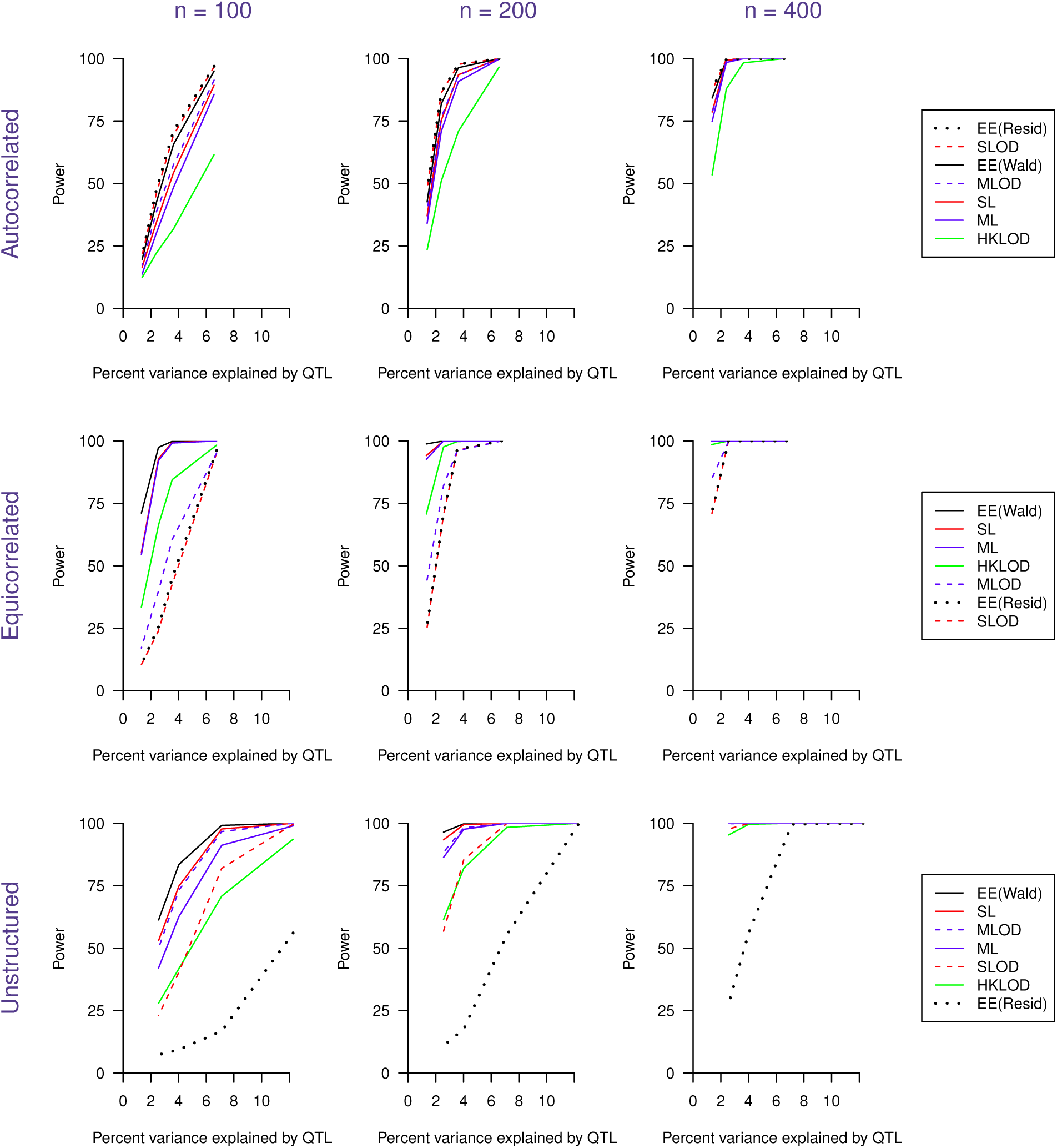
Power as a function of the percent phenotypic variance explained by a single QTL. The three columns correspond to sample sizes of *n* = 100, 200, and 400, respectively. The three rows correspond to the covariance structures (autocorrelated, equicorrelated, and unstructured). In the legends on the right, the methods are sorted by their overall average power for the corresponding covariance structure.

The precision of QTL mapping, measured by the root mean square error in the estimated QTL position, is displayed in Figure S1. Performance, in terms of precision, corresponds quite closely to the performance in terms of power: when power is high, the RMS error of the estimated QTL position is low, and vice versa.

A possible weakness of SLOD and MLOD approaches was that they do not make use of the function-valued form of the phenotypes. The methods may further suffer lower power in the case of noisy phenotypes. The methods proposed in this paper use smoothing to handle the measurement error. We repeated the same simulations in Kwak *et al.* (2014) with *n* = 200, adding independent, normally distributed errors (with standard deviation 1 or 2) at each time point.

The estimated power to detect the QTL as a function of heritability due to the QTL, for added noise with SD = 0,1,2 and the three different covariance structures, is shown in Figure 5. The power of the SLOD and MLOD were greatly affected by the introduction of noise. However, the SL and ML methods, which used both smoothing and dimension-reduction, work well in the presence of noise. The EE(Wald) method performed well in this case, as well. The EE(Residual) method did not work well compared to the other six methods. Overall, the EE(Wald) method continued to perform best.

**Figure 5:**
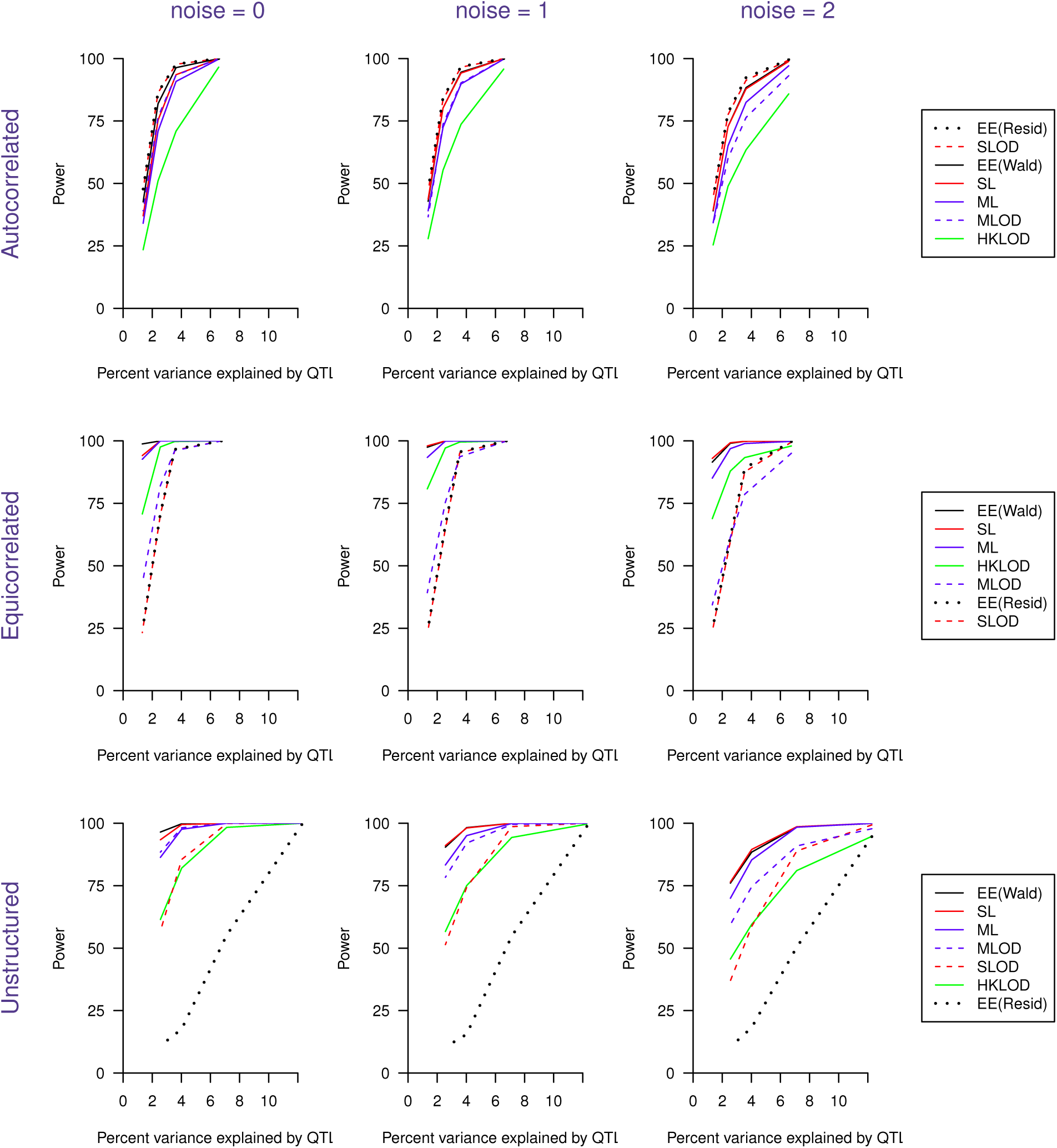
Power as a function of the percent phenotypic variance explained by a single QTL, with additional noise added to the phenotypes. The first column has no additional noise; the second and third columns have independent normally distributed noise added at each time point, with standard deviation 1 and 2, respectively. The percent variance explained by the QTL on the x-axis refers, in each case, to the variance explained in the case of no added noise. The three rows correspond to the covariance structures (autocorrelated, equicorrelated, and unstructured). In the legends on the right, the methods are sorted by their overall average power for the corresponding covariance structure.

Table 1 shows the average computation time for SLOD/MLOD, EE(Wald), EE(Residual), HKLOD and SL/ML methods. For the single-QTL simulations, the computation time for the SLOD/MLOD and HKLOD methods are similar, while the SL/ML methods are somewhat slower. This is because, in the simulation data set, the phenotype was measured at 10 time points. In the application, with 241 time points, the functional PCA based methods (HKLOD and SL/ML) are faster than the SLOD/MLOD methods. The EE(Wald) method requires considerably longer computation time; on the other hand, it provided the highest power to detect QTL.

**Table 1:**
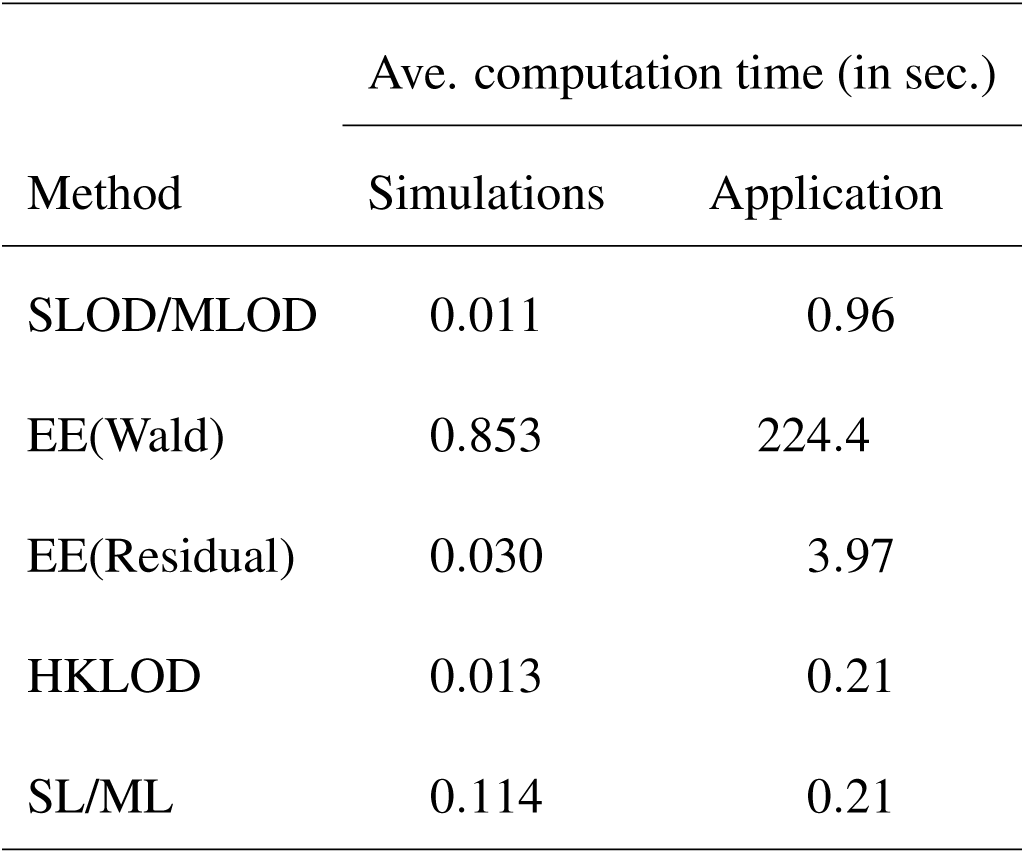
Average computation time for each method, in the single-QTL simulations and in the application.

### Multiple-QTL models

To investigate the performance of the penalized-HKLOD, penalized-SL and penalized-ML criteria in the context of multiple QTL, we used the same setting as Kwak *et al.* (2014). We simulated data from a three-QTL model modeled after that estimated, with the penalized-MLOD criterion, for the root tip angle data of Moore *et al.* (2013) considered in the Application section.

We assumed that the mean curve for the root tip angle phenotype followed a cubic polynomial, *y = a* + *bt* + *ct*^2^ + *dt*^3^, and assumed that the effect of each QTL also followed such a cubic polynomial. The four parameters for a given individual were drawn from a multivariate normal distribution with mean defined by the QTL genotypes and variance matrix estimated from the root tip angle data. Details on the parameter values used in the simulations appear in Kwak *et al.* (2014).

Normally distributed measurement error (with mean 0 and variance 1) was added to the phenotype at each time point for each individual. Phenotypes are taken at 241 equally spaced time points in the interval of 0 to 1. We considered two sample sizes: *n* = 162 (as in the Moore *et al.* (2013) data) and twice that, *n* = 324.

We performed 100 simulation replicates. For each replicate, we applied a stepwise model selection approach with each of the penalized-HKLOD, penalized-SL, penalized-ML, penalized-SLOD, and penalized-MLOD criteria. The simulation results are shown in Table 2.

**Table 2:**
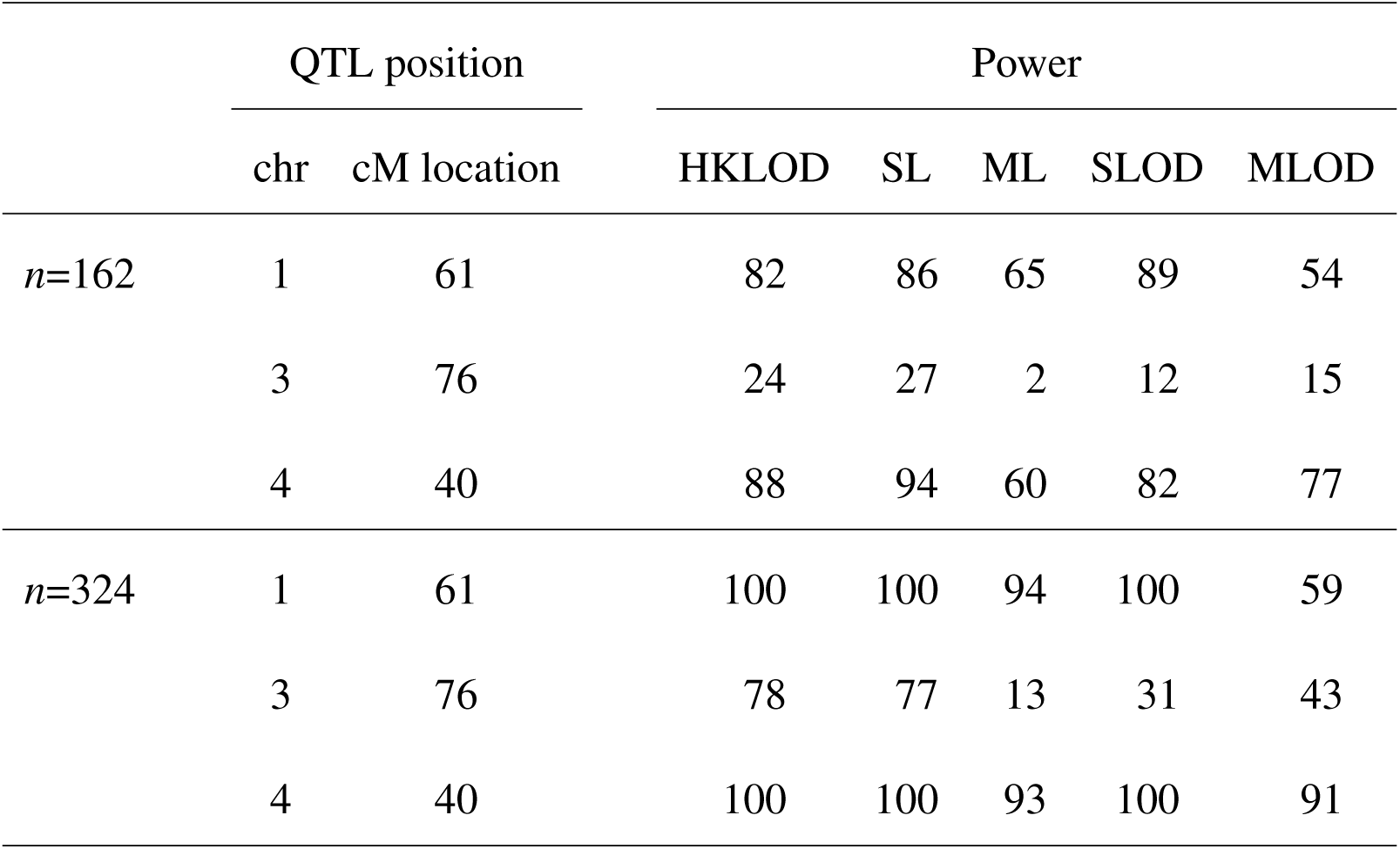
Power to detect QTL, for a three-QTL model modeled after the Moore *et al.* (2013) data.

The penalized-SL and penalized-HKLOD criteria had higher power to detect all three QTL. In particular, the power to detect the QTL on chromosome 3 was greatly increased, in comparison to the penalized-SLOD and penalized-MLOD criteria.

## Discussion

We have described two techniques for improving the regression-based methods of Kwak *et al.* (2014) for QTL mapping with function-valued phenotypes: smoothing and dimensional-reduction. Smoothing leads to better performance in the case of noisy phenotype measurements (Li and Sillanpää 2013), and dimension-reduction improves power. The particular methods we used (smoothing via B-splines, and dimensional reduction via functional principal component analysis) are not the only possibilities, but they are natural choices widely used in functional data analysis (Ramsay and Silverman 2005).

Following smoothing and dimensional-reduction, we applied QTL analysis to the small number of derived traits, either by analyzing the traits individually and then combining the log likelihoods, as in Kwak *et al.* (2014), or by applying the multivariate QTL mapping method of Knott and Haley (2000). The latter approach cannot be applied directly to the original phenotypes, due to the large number of time points at which the traits were measured, but it can work well with the dimension-reduced derived traits.

Key advantages of our proposed methods include speed of computation and the ability to consider multiple-QTL models. The EE(Wald) method of Xiong *et al.* (2011), based on estimating equations, was seen to be most powerful for QTL detection in our simulation study, but it is orders of magnitude slower and has not yet been implemented for multiple-QTL models.

Many other methods have been developed for QTL mapping with function-valued traits. However, most focus on single-QTL models (e.g., Ma *et al.* 2002; Yang *et al.* 2009; Yap *et al.* 2009). Bayesian methods for multiple-QTL mapping with function-valued traits have been proposed (Min *et al.* 2011; Sillanpää *et al.* 2012; Li and Sillanpää 2013), but these methods are computationally intensive, and software is not available.

In considering multiple-QTL models, we have focused on strictly additive QTL. Manichaikul *et al.* (2009) extended the work of Broman and Speed (2002) by considering pairwise interactions among QTL. Our approaches may be similarly extended to handle interactions.

The enormous recent growth in capabilities for high-throughput phenotyping, including images and time series, particularly in plants (see, for example, Cabrera-Bosquet *et al.* 2012; Araus and Cairns 2014; Ghanem *et al.* 2015), is accompanied by a growth in interest in the genetic analysis such phenotype data. Speed of computation will be particularly important in the analysis of such high-dimensional data, as will the joint consideration of multiple loci. The methods we have proposed can meet many of these challenges.

Our efforts on this problem were inspired by the data of Moore *et al.* (2013), in which the phenotype was measured at a large number of time points. While our approaches do not require such a high density of time points, we expect that the use of smoothing and functional PCA will be most suitable in the case of at least 8–10 time points. In the case of a smaller number of time points, we would recommend the direct use of multivariate QTL analysis, or of the SLOD and MLOD methods of Kwak *et al.* (2014).

We implemented our methods as a package, funqtl, for the general statistical software R (R Core Team 2015). Our package makes use of the fda package (Ramsay *et al.* 2014) for smoothing and functional PCA. It is available at https://github.com/ikwak2/funqtl.

## Acknowledgments

The authors thank Nathan Miller and two anonymous reviewers for suggestions to improve the manuscript. This work was supported in part by grant IOS-1031416 from the National Science Foundation Plant Genome Research Program to E.P.S. and by National Institutes of Health grant R01GM074244 to K.W.B.

